# Overexpression of the Rieske FeS protein of the Cytochrome *b*_6_*f* complex increases C_4_ photosynthesis

**DOI:** 10.1101/574897

**Authors:** Maria Ermakova, Patricia E. Lopez-Calcagno, Christine A. Raines, Robert T. Furbank, Susanne von Caemmerer

## Abstract

C_4_ plants contribute 20% to the global primary productivity despite representing only 4% of higher plant species. Their CO_2_ concentrating mechanism operating between mesophyll and bundle sheath cells increases CO_2_ partial pressure at the site of Rubisco and hence photosynthetic efficiency. Electron transport chains in both cell types supply ATP and NADPH for C_4_ photosynthesis. Since Cytochrome *b*_6_*f* is a key point of control of electron transport in C_3_ plants, we constitutively overexpressed the Rieske FeS subunit in *Setaria viridis* to study the effects on C_4_ photosynthesis. Rieske FeS overexpression resulted in a higher content of Cytochrome *b*_6_*f* in both mesophyll and bundle sheath cells without marked changes in abundances of other photosynthetic complexes and Rubisco. Plants with higher Cytochrome *b*_6_*f* abundance showed better light conversion efficiency in both Photosystems and could generate higher proton-motive force across the thylakoid membrane. Rieske FeS abundance correlated with CO_2_ assimilation rate and plants with a 10% increase in Rieske FeS content showed a 10% increase in CO_2_ assimilation rate at ambient and saturating CO_2_ and high light. Our results demonstrate that Cytochrome *b*_6_*f* controls the rate of electron transport in C_4_ plants and that removing electron transport limitations can increase the rate of C_4_ photosynthesis.

## Introduction

Crop yield gains achieved during the Green Revolution by conventional plant breeding were not based on photosynthetic traits^1^ and the theoretical maximum yield of the light conversion in crops is yet to be achieved^2^. Increasing photosynthetic efficiency by targeted genetic manipulation could potentially double the yield of crop plants^3^. Electron transfer reactions of photosynthesis (also called light reactions) supply ATP and NADPH essential for CO_2_ assimilation and therefore are a target for improvement^4^. It has been demonstrated in C_3_ plants that facilitating electron transport by overexpressing the components of electron transfer chain can result in higher assimilation rates^5, 6^. As C_4_ plants play a key role in world agriculture with maize and sorghum being major contributors to world food production and sugarcane, miscanthus and switchgrass being major plant sources of bioenergy^7^, improvement of electron transport reactions could further increase the rates of C_4_ photosynthesis and yield^8, 9^.

In C_3_ plants the electron transport chain is localised to the thylakoids membranes of mesophyll cells and consists of four major protein complexes: Photosystem II (PSII), Cytochrome *b*_6_*f* (cyt*b*_6_*f*), Photosystem I (PSI) and ATP synthase. The first three complexes sustain linear electron flow from the water-oxidizing complex of PSII to NADP^+^, the terminal acceptor of PSI, which is accompanied by acidification of the internal compartments of thylakoids (lumen). Transmembrane difference in electrochemical potentials of protons, or proton-motive force (*pmf*), serves as the driving force for ATP synthesis.

Cyt*b*_6_*f* oxidises plastoquinol reduced by PSII and reduces plastocyanin, which then diffuses to PSI; plastoquinol oxidation is a rate-limiting step in the intersystem chain^10^. As a result of Q-cycle operating between the two binding sites of cyt*b*_6_*f*, two protons are translocated from the stroma to the lumen per one electron^11^. Once pools of inorganic phosphate are exhausted, ATP synthesis slows down and causes a build-up of the *pmf* across the membrane^12^. Both components of *pmf*, the pH gradient (ΔpH) and the electrochemical gradient (ΔΨ), are equally capable of driving ATP synthesis^13^. However, the pH component has a major regulatory effect on the electron transport chain and at pH<6 causes slowing down of the plastoquinol oxidation^14^ and down-regulation of PSII activity via non-photochemical quenching (NPQ). NPQ is a mechanism that trigs the attenuation of PSII activity by the dissipation of excess light energy in the form of heat in the light-harvesting antenna of PSII (LHCII)^15^. Fast, pH-dependent NPQ component in higher plants, q_E_, is regulated by the PHOTOSYSTEM II SUBUNIT S (PsbS) protein^16^ and xanthophyll cycle^17^.

Cyt*b*_6_*f* forms a homodimer where each monomer consists of eight subunits: major subunits, Rieske FeS protein (PetC), cyt *b*_6_ (PetB), cyt *f* (PetA) and subunit IV (PetD), and minor subunits, PetG, PetL, PetM and PetN^18^. There is an increasing amount of evidence that the amount of Rieske FeS protein, one of the two nuclear-coded subunits along with PetM, regulates the abundance of cyt*b*_6_*f*^6, 19–24^. Transgenic *Arabidopsis thaliana* plants overexpressing Rieske FeS also showed an increase in the amounts of other cyt*b*_6_*f* subunits, positive effects PSII electron transport rate and CO_2_ assimilation rate and decreased NPQ^6^. Studies on transgenic tobacco plants indicate that cyt*b*_6_*f* determines the rate of electron transport through the electron transport chain and concomitantly the CO_2_ assimilation rate^19–23^.

C_4_ photosynthesis is a biochemical CO_2_ concentrating pathway operating between mesophyll (M) and bundle sheath (BS) cells and there are three biochemical C_4_ subtypes^25^. PEP carboxylase (PEPC) catalyses primary carbon fixation in the cytoplasm of mesophyll cells into C_4_ acids. In plants of NADP-ME subtype, like maize, sorghum and setaria, malate diffuses to the bundle sheath cells where it is decarboxylated inside chloroplasts to provide CO_2_ for ribulose bisphosphate carboxylase oxygenase (Rubisco). Pyruvate resulting from malate decarboxylation diffuses back to mesophyll cells where it is regenerated into PEP by pyruvate ortophosphate dikinase. C_4_ species with NADP-ME biochemistry require a minimum of 1 NADPH and 2 ATP in mesophyll cells and 1 NADPH and 3 ATP in bundle sheath cells per one CO_2_ fixed^8^. The components of mesophyll electron transport chain are very similar to those described above for C_3_ plants but bundle sheath cells of NADP-ME C_4_ species are effectively supplied with NADPH via malate coming from the mesophyll cells and therefore are more specialised for ATP production. Bundle sheath cells of NADP-ME species usually have little or no PSII activity and operate active cyclic electron flow (CEF) leading to the formation of *pmf* but not to NADP^+^ reduction^26^. There are two pathways for CEF: one via PROTON GRADIENT REGULATION 5 protein (PGR5), cytb6f and PSI, and another one via chloroplastic NAD(P)H:Quinone oxidoreductase 1-like complex, cyt*b*_6_*f* and PSI^27^.

Since cyt*b*_6_*f* is a component of electron transport chain in both mesophyll and bundle sheath cells, we used *Setaria viridis*, a model NAPD-ME-type C_4_, plant to study effects of constitutive Rieske FeS overexpression on C_4_ photosynthesis. We demonstrate that in both cell types increased abundance of Rieske FeS results in higher content of cyt*b*_6_*f* and allows higher photosynthesis rates without notable changes of Rubisco and chlorophyll content. Our results indicate that in C_4_ plants electron transport is one of the limitations for CO_2_ assimilation, particularly at high light and non-limiting CO_2_ concentrations, and it is under cyt*b*_6_*f* control.

## Results

### Generation and selection of transgenic *S.viridis* plants with Rieske FeS overexpression

For Rieske FeS overexpression, the coding sequence of PetC gene from *Brachypodium distachyon* (*Bd*PetC) was codon-optimized for the Golden Gate cloning system and assembled into two constructs (230 and 231) under the control of the maize ubiquitin promoter (Fig. S1). The constructs were transformed into *S.viridis* using stable agrobacterium-mediated transformation. Eleven T_0_ transgenic plants were selected on the basis of hygromicin resistance and a subset of nine lines was analysed for Rieske FeS protein abundance, the presence of the *Bd*PetC transcript and insertion numbers (Fig. 1a). Plants that went through the transformation process but tested negative for the insertion were used as control for T_0_ plants and T_1_ progeny. Rieske FeS protein abundance per leaf area was highly variable between transgenic and control plants in the T_0_ generation (Fig. 1a). Therefore three lines, 230(4), 231(3) and 231(6), were selected for further analysis of the T_1_ progeny based on the presence of the *Bd*PetC transcript (Fig. 1a).

**Fig. 1.**
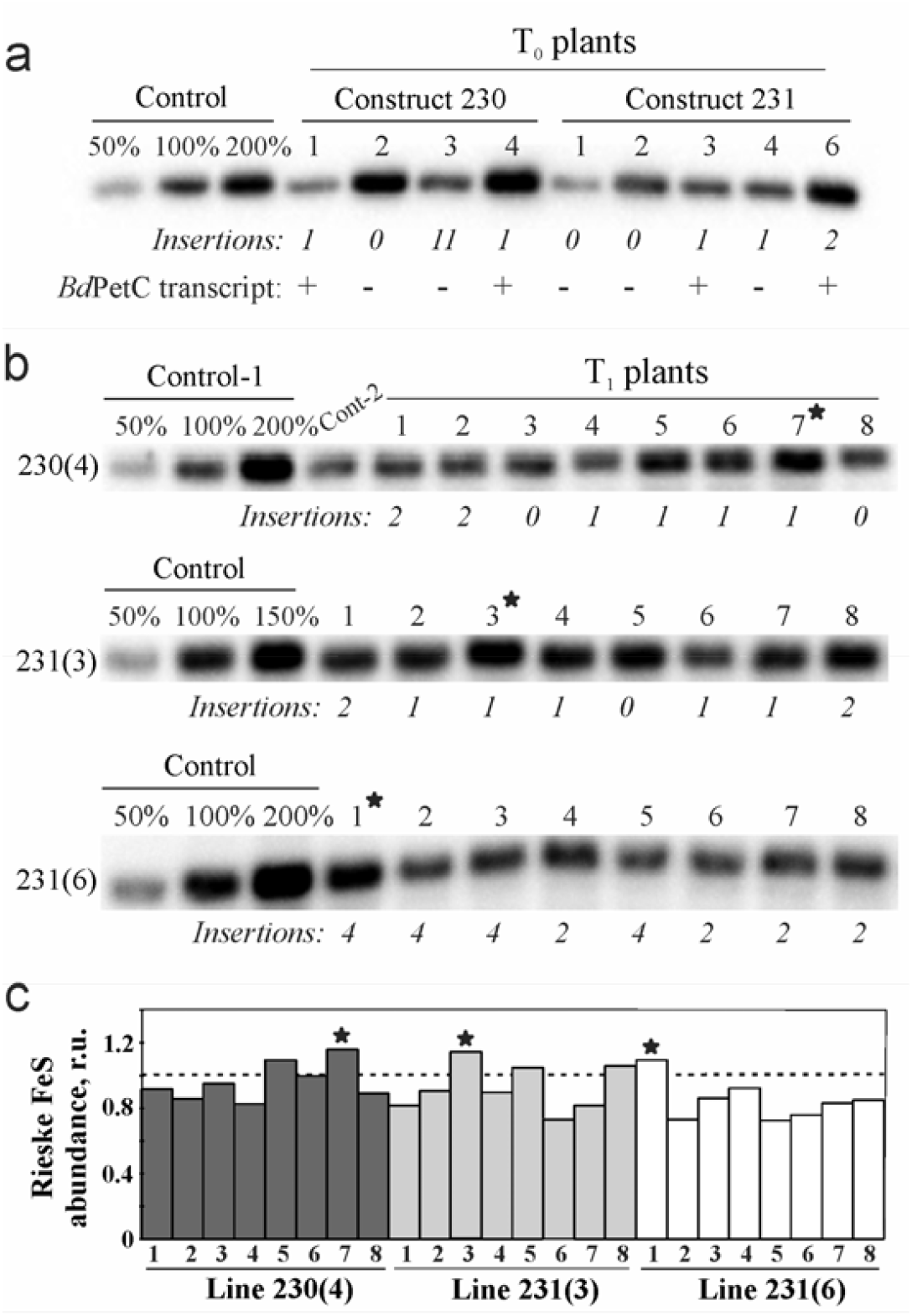
Selection of the transgenic *S.viridis* plants with Rieske FeS overexpression by immunodetection of Rieske FeS protein abundance on leaf area basis. **a**. T_0_ lines selected based on hygromycin resistance after transformation with constructs 230 and 231; insertion numbers indicate copy numbers of the hygromycin phosphotransferase gene; (+), transcript of the recombinant PetC from *B.distachyon* (*Bd*PetC) is present; (−) *Bd*PetC transcript is not present. **b**. T_1_ plants of three independent lines and their insertion numbers. **c**. Quantitative estimate of Rieske FeS abundance in T_1_ plants from the immunoblots shown in (b) relative to control plants (=1). The plants that went through the transformation process and tested negative for the hygromycin phosphotransferase gene were used as control plants in the T_0_ and T_1_ generations. Asterisks indicate plants selected for further analysis in the T_2_ generation.

In the T_1_ generation, a quantitative estimate of the changes in Rieske FeS levels from the immunoblots showed that a number of transgenic plants contained increased amounts of Rieske FeS protein per leaf area compared to control plants with a maximum increase of 10-15% in 230(4)-7, 231(3)-3 and 231(6)-1 plants (Fig. 1b, Fig. 1c). Seeds from those plants were collected and used for the analysis of their T_2_ progeny. Interestingly, analysis of the *Bd*PetC expression in the T_1_ generation of line 230(4) (Fig. S2) demonstrated that, although homozygous plants had about 2-fold higher transcript abundance compared to heterozygous plants, they showed no increase of Rieske FeS protein level. Similar effect was observed in T_1_ plants of line 231(3) and consequently also in the T_2_ progeny of line 231(3)-3: only some heterozygous plants had increased Rieske FeS protein abundance (Fig. 2b, Fig. S3). Moreover, Rieske FeS protein overexpression could only be detected when plants were grown at a low irradiance of 200 μmol m^−2^ s^−1^ suggesting a post-translational and light-dependent regulation of Rieske FeS abundance in *S.viridis*.

**Fig. 2.**
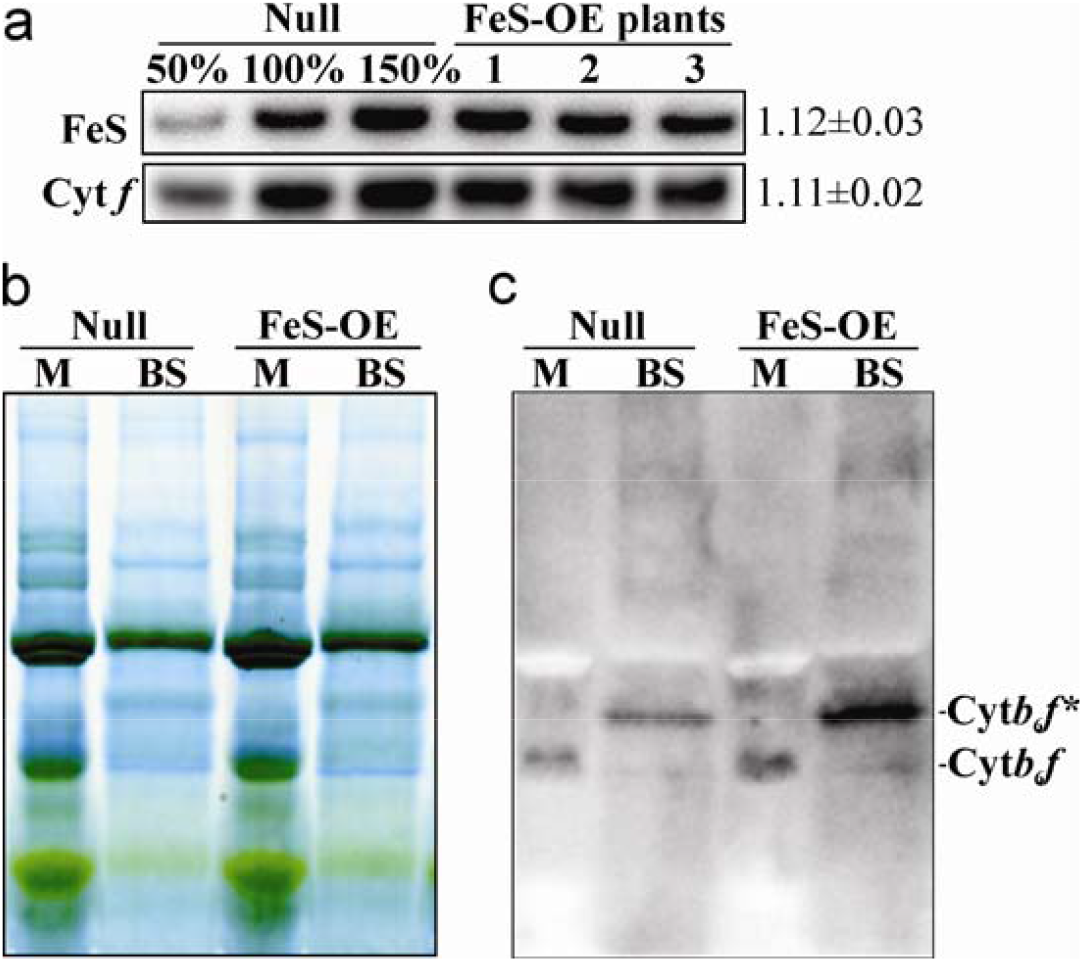
Relative abundance of Cytochrome *b*_6_*f* complex in plants with Rieske FeS overexpression (FeS-OE) and Null plants. **a**. Immunodetection of the Rieske FeS and Cytochrome *f* subunits and quantification of the relative protein abundance per leaf area in T_2_ plants of line 230(4)-7 compared to Null (=1). Mean ± SE, n=3. **b**. Blue Native gel electrophoresis of the thylakoid protein complexes isolated from mesophyll (M) and bundle sheath (BS) cells; 10 μg of chlorophyll (*a+b*) loaded in each lane. **c.** Immunodetection of Cytochrome *b*_6_*f* complex from the Blue-Native gel with Rieske FeS antibodies; Cyt*b*_6_*f* and Cyt*b*_6_*f*\* - two distinct forms of the complex detected.

Transgenic plants with higher Rieske FeS protein abundance (FeS-OE hereafter) could be identified using pulse modulated chlorophyll fluorescence measurements of photochemical and non-photochemical processes at the incident light intensity with the MultispeQ (Table 1). FeS-OE plants demonstrated significantly higher quantum yield of PSII (φPSII), a parameter estimating a proportion of the light absorbed by chlorophyll associated with PSII that is used in photochemistry, and a lower quantum yield of NPQ (φNPQ), i.e. a proportion of absorbed light actively dissipated in the PSII antennae. Electron transport parameters measured in transgenic plants harbouring the transgene but not showing an increase in Rieske FeS abundance (“No FeS-OE” in Table 1) did not differ from Nulls. Other parameters such as relative chlorophyll content (SPAD) and leaf thickness did not differ between the three groups (Table 1).

**Table 1.** Photosynthetic electron fluxes and leaf parameters in transgenic and Null plants. “No FeS-OE”, T_2_ plants of line 230(4)-7 with the Null levels of Rieske FeS protein; “FeS-OE”, T_2_ plants of line 230(4)-7with higher abundance of Rieske FeS per leaf area. MultispeQ measurements were performed at the incident irradiance of 200 μmol m^−2^ s^−1^, mean ± SE, n=16-20 (3-4 measurements per plant): φPSII, quantum yield of PSII; φNPQ, quantum yield of non-photochemical quenching; φNO, quantum yield of non-regulatory energy dissipation. Chlorophyll and Rubisco measurements are mean ± SE, n=3. Asterisks indicate statistically significant differences between transgenic and Null plants (*P*<0.05).

| Parameter | Null | No FeS-OE | FeS-OE |
| --- | --- | --- | --- |
| Relative chlorophyll (SPAD) | 50.79±1.60 | 51.56±1.61 | 51.78±2.06 |
| Leaf thickness, mm | 0.76±0.17 | 0.74±0.12 | 0.72±0.11 |
| $\phi\text{PSII}$ | 0.598±0.010 | 0.597±0.004 | 0.624±0.004* |
| $\phi\text{NPQ}$ | 0.205±0.011 | 0.200±0.005 | 0.182±0.003* |
| $\phi\text{NO}$ | 0.197±0.002 | 0.201±0.001 | 0.194±0.002 |
| Chlorophyll (a+b), g m<sup>-2</sup> | 0.39±0.06 | | 0.38±0.05 |
| Chlorophyll a/b | 5.05±0.13 | | 5.10±0.10 |
| Rubisco, g m<sup>-2</sup> | 0.35±0.03 | | 0.34±0.02 |

### Rieske FeS overexpression results in higher abundance of the Cytochrome *b*_6_*f* complex

To see if the higher level of Rieske FeS protein resulted in increased abundance of the whole cyt*b*6*f* complex, abundance of Rieske FeS and cytochrome *f* subunits of cyt*b*6*f* was analysed on leaf area basis in the T_2_ progeny of 230(4)-7 using specific antibodies (Fig. 2a). Both cyt*b*_6_*f* subunits were more abundant in FeS-OE plants relative to Null with a significant increase of 1.12±0.03 and 1.11±0.02-fold, respectively (mean ± SE, n=3).

Thylakoid protein complexes from the bundle sheath and mesophyll cells of FeS-OE and Null plants were separated using Blue Native gel electrophoresis. No major changes were found between transgenic and Null plants in composition of complexes within the thylakoid membranes of each cell type (Fig. 2b). Thylakoid protein complexes were probed with Rieske FeS antibodies to obtain the relative abundance of cyt*b*_6_*f* complex. Whilst one band matching the cyt*b*_6_*f* band reported previously^28^ was detected In the mesophyll thylakoids, in the bundle sheath thylakoids two bands were recognized by the Rieske FeS antibodies: a minor one with a similar molecular weight to the mesophyll cyt*b*6*f* and a major one with a slightly higher molecular weight (cyt*b*_6_*f*\*) (Fig. 2c). Relative abundance of cyt*b*_6_*f* was higher in the mesophyll thylakoids and cyt*b*_6_*f*\* was more abundant in the bundle sheath thylakoids of FeS-OE plants compared to Nulls (Fig. 2c).

### Rieske FeS overexpression increases CO_2_ assimilation rate and quantum yield of Photosystem II

CO_2_ assimilation rate was measured in the T_1_ plants of lines 230(4), 231(3) and 231(6) and the T_2_ progenies of lines 230(4)-7, 231(3)-3 and 231(6)-1 at 1500 μmol quanta m^−2^ s^−1^ and CO_2_ of 400 ppm and plotted against the relative abundance of Rieske FeS protein (Fig. 3). All three lines showed a positive correlation between Rieske FeS abundance and CO_2_ assimilation rate, suggesting that higher abundance of cyt*b*_6_*f* supported higher rates of C_4_ photosynthesis.

**Fig. 3.**
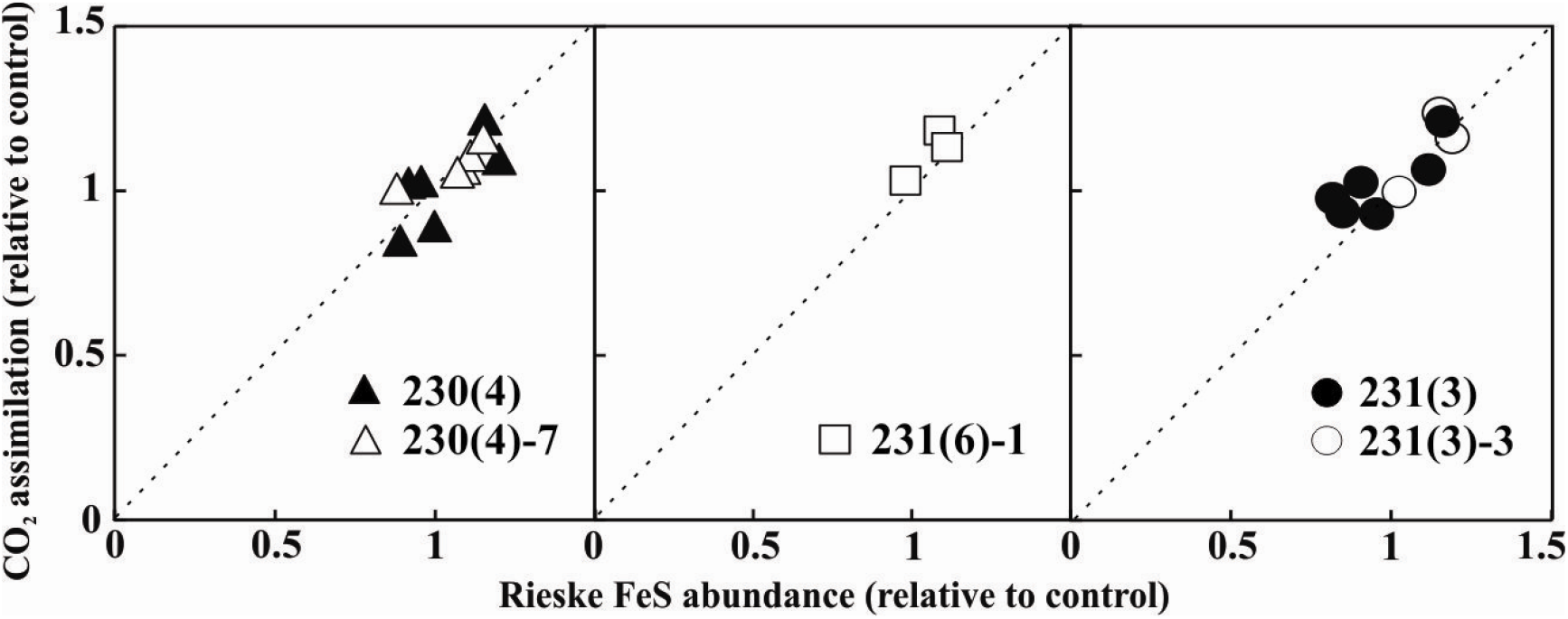
Relationships of Rieske FeS abundance and CO_2_ assimilation rate in T_1_ (filled symbols) and T_2_ (open symbols) plants of three independent lines relative to control plants. Rieske FeS abundance was quantified from the immunoblots shown on the Fig. 1b, Fig. 2b, Fig. S3. Steady-state CO_2_ assimilation rates were measured at 1500 μmol m^−2^ s^−1^ and CO_2_ partial pressure of 400 ppm. The plants that went through the transformation process and tested negative for the hygromycin phosphotransferase gene were used as control for T_1_ plants and Null segregates were used as control for T_2_ plants. Average rates of CO_2_ assimilation measured from control plants were 26.47 μmol m^−2^ s^−1^ for T_1_ plants and 32.65 μmol m^−2^ s^−1^ for T_2_ plants.

To examine the effect of Rieske FeS overexpression on CO_2_ assimilation rate in more detail, the CO_2_ response of assimilation was examined at a constant irradiance of 1500 μmol m^−2^ s^−1^ (Fig. 4a) in the T_2_ progenies of lines 230(4)-7 and 231(3)-3. At lower intercellular CO_2_ partial pressure no differences in assimilation rates were found between two FeS-OE lines and Null plants (Fig. 4a, Fig. S4). However, at C_i_ above 150 μbar FeS-OE plants had higher CO_2_ assimilation rates with a significant increase at the C_i_ above 250 μbar. Maximum CO_2_ assimilation rates reached by FeS-OE plants of line 230(4) and Null plants at high C_i_ were 41.30±0.89 and 38.14±0. 818 μmol m^−2^ s^−1^ (mean ± SE, n=4), respectively, indicating about a 10% increase in photosynthesis in FeS-OE plants (Fig. 4a). FeS-OE plants of the line 231(3) also demonstrated about 10% increase of CO_2_ assimilation rates at ambient and high CO_2_ (Fig. S4). CO_2_ response of φPSII measured in FeS-OE and Null plants showed similar trends: no difference at low C_i_ but significantly higher φPSII in FeS-OE plants at C_i_ above 250 μbar (Fig. 4b). Stomatal conductance in FeS-OE plants estimated from the gas-exchange measurements was higher at Ci above 250 μbar but not significantly (0.05<*P*<0.15) (Fig. 4c).

**Fig. 4.**
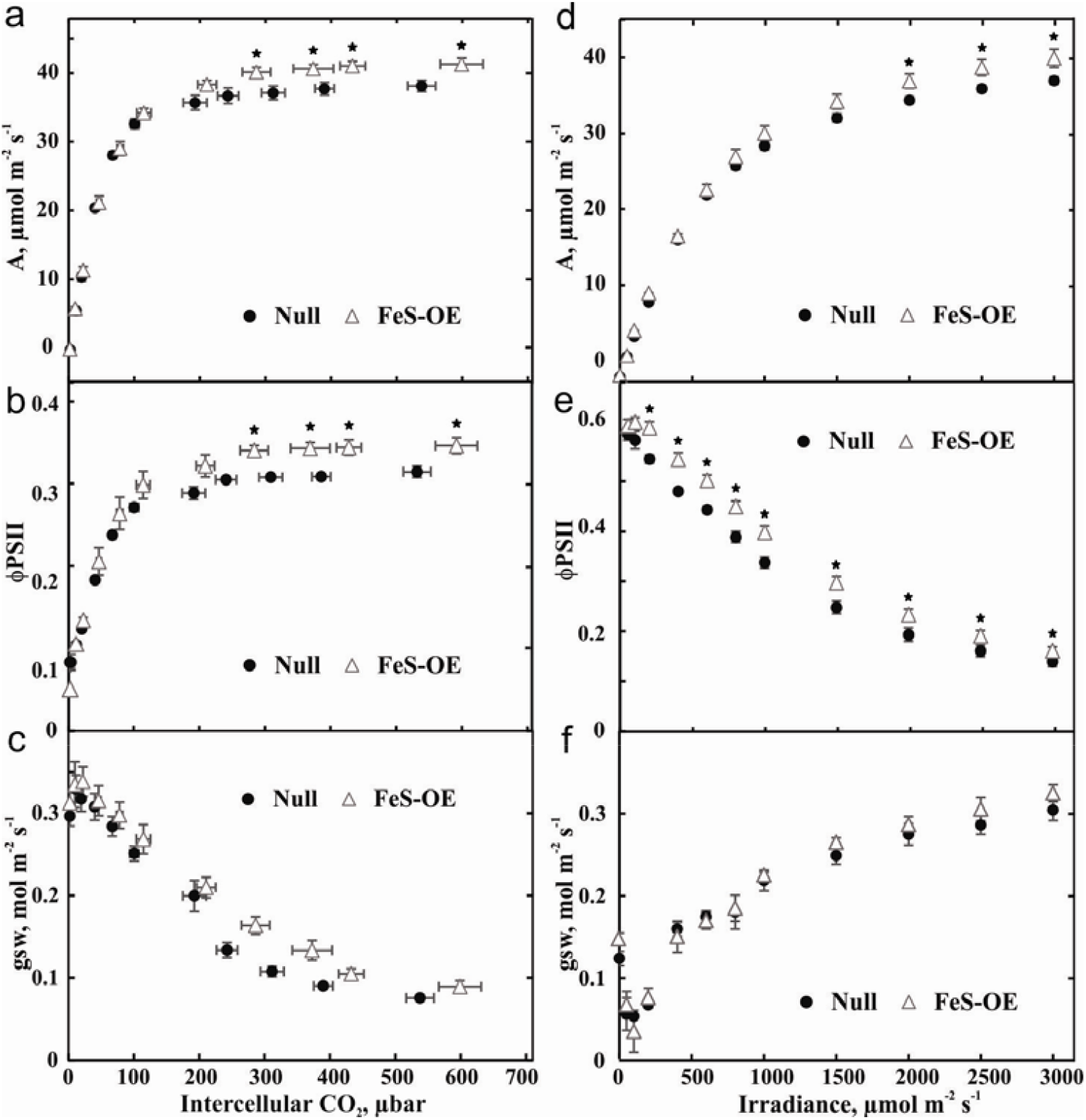
Gas-exchange and fluorescence analysis of plants with Rieske FeS overexpression (FeS-OE) and Null plants. **a. b. c.** CO_2_ response of assimilation rate (A), quantum yield of Photosystem II (φPSII) and stomatal conductance (gsw) at 1500 μmol m^−2^ s^−1^. **d. e. f.** Light response of A, φPSII and gsw at CO_2_ partial pressure of 400 ppm. Mean ± SE, n=4. Asterisks indicate statistically significant differences between FeS-OE and Null plants (*P*<0.05). Results are for the T_2_ progeny of line 230(4)-7, results for the T_2_ progeny of line 231(3)-3 are shown in Fig. S4.

The light response of CO_2_ assimilation was examined at a constant CO_2_ of 400 ppm. Assimilation rates did not differ between FeS-OE plants of line 230(4)-7 and Null plants at the range of irradiances below 1500 μmol m^−2^ s^−1^ (Fig. 4d). However at light levels above 1500 μmol m^−2^ s^−1^, FeS-OE plants had significantly higher assimilation rates compared to Nulls with a maximum increase of about 8%. φPSII measured concurrently was significantly higher in FeS-OE plants compared to Null plants over the range of irradiances above 200 μmol m^−2^ s^−1^ (Fig. 4e) whilst stomatal conductance did not differ in transgenic plants independently on irradiance (Fig. 4f).

### Rieske FeS overexpression plants have lower NPQ and generate more proton-motive force

NPQ as well as photochemical and non-photochemical yields of PSI at different irradiances were analysed in FeS-OE plants of line 230(4)-7 and Null plants with the Dual-PAM-100. In agreement with the higher φPSII, FeS-OE plants also had significantly lower NPQ at the irradiances above 220 μmol m^−2^ s^−1^ (Fig. 5a). The effective quantum yield of PSI (φPSI) in FeS-OE plants was significantly higher compared to Null plants at the irradiances above 57 μmol m^−2^ s^−1^ (Fig. 5b). φND, a non-photochemical loss due to the oxidised primary donor of PSI, was significantly lower in FeS-OE plants compared to Nulls at the range of irradiances between 57 and 220 μmol m^−2^ s^−1^ (Fig. 5c). φNA, a non-photochemical loss due to the reduced PSI acceptor, did not differ significantly between FeS-OE and Null plants but FeS-OE showed a tendency to lower φNA at high irradiance (0.1>*P*>0.05) (Fig. 5d).

**Fig. 5.**
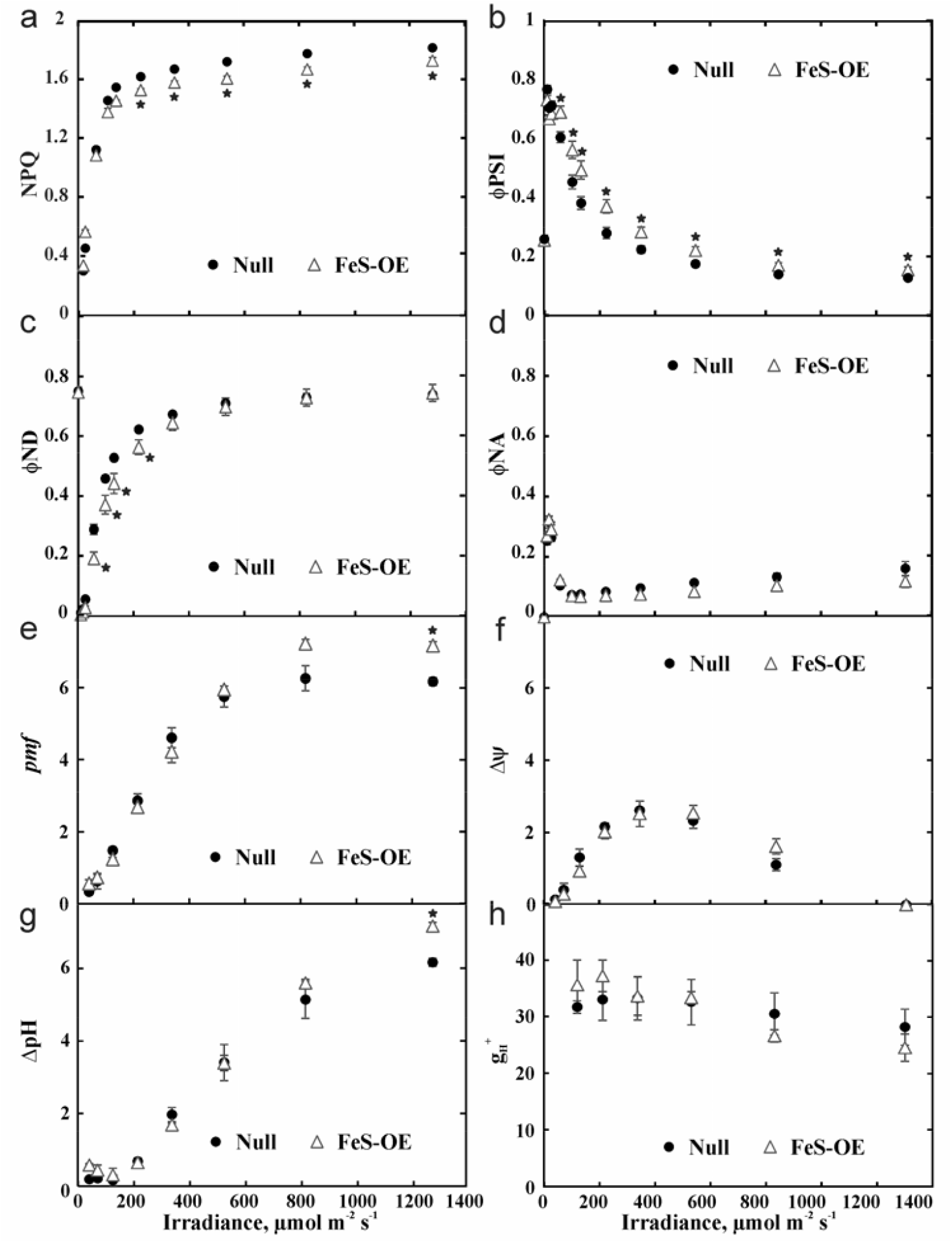
Light response of photosynthetic parameters in plants with Rieske FeS overexpression (FeS-OE) and Null plants. **a.** NPQ, non-photochemical quenching. **b.** φPSI, quantum yield of PSI. **c.** ND, non-photochemical loss due to the oxidised primary donor of PSI. **d.** φND, non-photochemical loss due to the reduced PSI acceptor. **e.** *pmf*, proton-motive force, estimated from the electrochromic shift signal. **f.** Δψ, electrochemical gradient component of *pmf*. **g.** ΔpH, proton gradient component of *pmf*. **h.** g_H_^+^, proton conductivity of the thylakoid membrane. Results are for the T_2_ progeny of line 230(4)-7. Mean ± SE, n=3. Asterisks indicate statistically significant differences between FeS-OE and Null plants (*P*<0.05).

Dark-interval relaxation of the electrochromic shift signal recorded to explore the generation of *pmf* in leaves at various irradiances. Total *pmf* was significantly higher in FeS-OE plants at 1287 μmol m^−2^ s^−1^ (Fig. 5e). Decomposition of *pmf* into electrochemical gradient (ΔΨ) and proton gradient (ΔpH) components revealed that the higher *pmf* at 1287 μmol m^−2^ s^−1^ was due to the higher ΔpH (Fig. 5f, Fig. 5g). Fitting of the first-order ECS relaxation kinetics showed that there was no difference in thylakoid proton conductance (g_H_^+^) between FeS-OE and Null plants irrespective of irradiance (Fig. 5h).

Light-induced changes of photosynthetic parameters were studied on dark-adapted leaves over the course of illumination with red actinic light of 220 μmol m^−2^ s^−1^ (Fig. 6). During the first three minutes of illumination, FeS-OE plants had significantly lower NPQ indicating a slower induction of q_E_ compared to Null plants whilst no difference in NPQ relaxation kinetics was observed in darkness. Higher φPSII during the first minutes of irradiance correlated with the lower NPQ in FeS-OE plants. φPSI was higher in FeS-OE plants over the whole course of illumination compared to Null plants (Fig. 6).

**Fig. 6.**
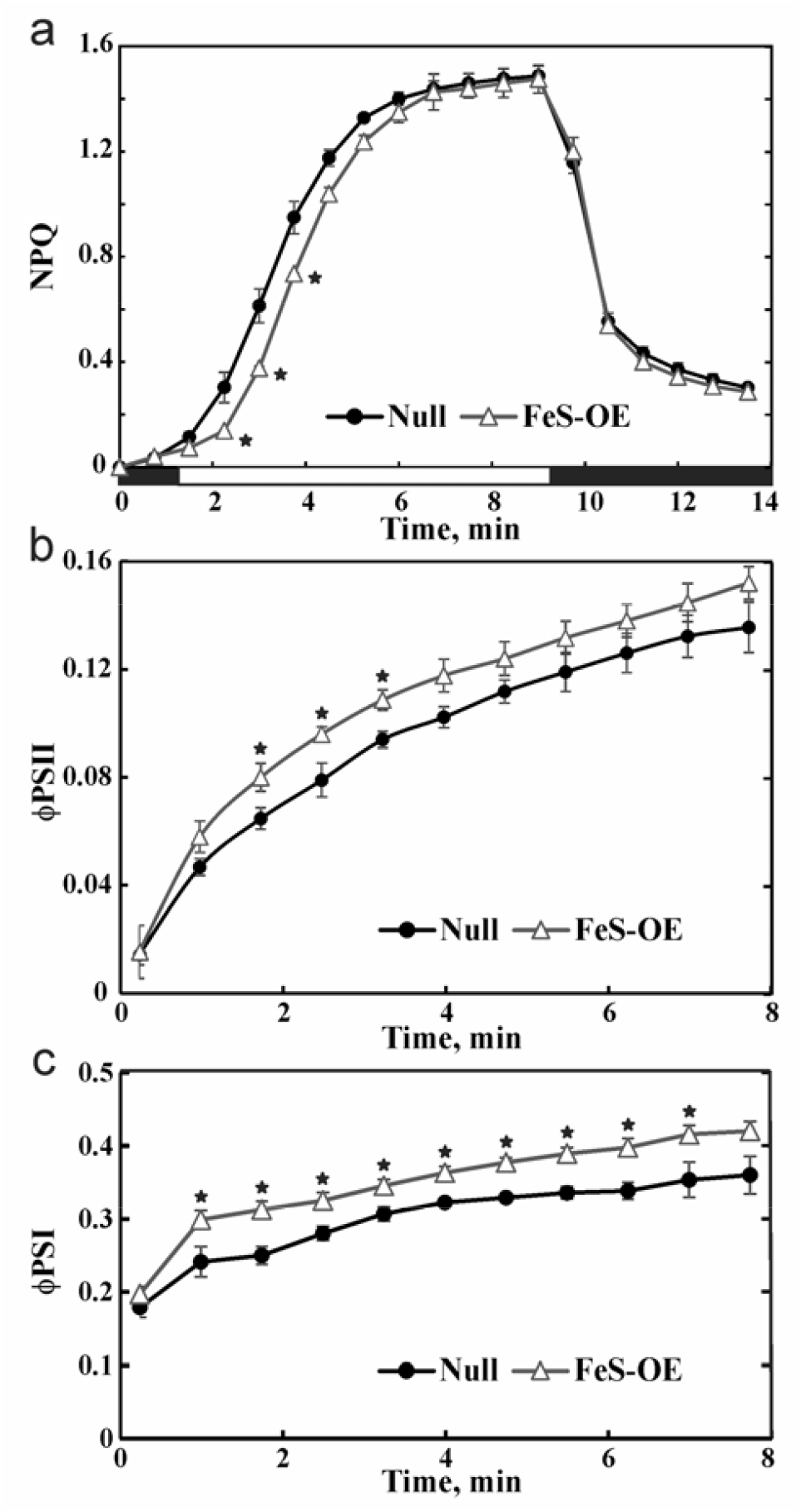
Photosynthetic parameters of plants with Rieske FeS overexpression (FeS-OE) and Null plants during dark/light transitions measured on dark-adapted leaves. **a.** Induction and relaxation of NPQ during dark-light-dark shift: black bars, darkness; white bar, red actinic light of 220 μmol m^−2^ s^−1^. **b. c.** Dynamic changes of φPSII and φPSI, quantum yields of Photosystem II and Photosystem I, during first minutes of illumination with red actinic light of 220 μmol m^−2^ s^−1^. Results are for T_2_ plants of line 230(4)-7. Mean ± SE, n=3. Asterisks indicate statistically significant differences between FeS-OE and Null plants (*P*<0.05).

### Levels of other photosynthetic proteins are unchanged in FeS-OE plants

Immunoblotting with specific antibodies against representative subunits of photosynthetic thylakoid complexes suggested that there were no significant differences in abundances of PSI, PSII, ATP synthase, NDH and light-harvesting complexes of PSI and PSII between FeS-OE and Null plants on the leaf level (Fig. 7). In addition, no changes in PsbS abundance were detected. Abundances of Rubisco large subunit (RbcL) and PEPC in FeS-OE plants were also similar to the levels detected in Null plants (Fig. 6). In line with the immunoblotting results, amounts of Rubisco active centers in FeS-OE and Null plants, estimated by [^14^C] carboxyarabinitol bisphosphate binding assay, did not differ and neither did leaf chlorophyll (*a+b*) content and chlorophyll *a/b* ratio (Table 1).

**Fig. 7.**
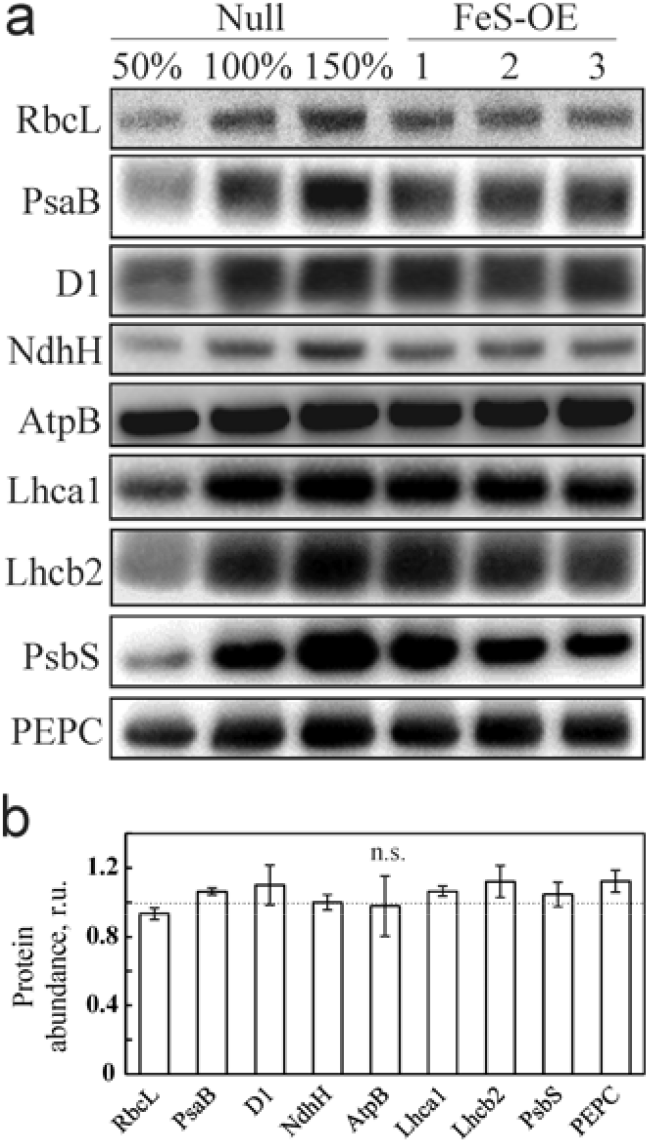
Immunodetection of photosynthetic proteins in T_2_ plants of line 230(4)-7 with Rieske FeS overexpression (FeS-OE) and Null plants on leaf area basis. **a.** Western blots with antibodies against RbcL (large subunit of Rubisco), PsaB (Photosystem I), D1 (Photosystem II), NdhH (NDH complex), AtpB (ATP synthase), Lhca1 (light-harvesting complex of Photosystem I), Lhcb2 (light-harvesting complex of Photosystem II), PsbS and PEPC. **b.** Quantification of protein abundances in FeS-OE plants relative to Null (=1). Mean ± SE, n=3. No statistically significant differences detected.

## Discussion

C_4_ plants already have high photosynthetic rates when compared to C_3_ plants which is achieved by operating a biochemical carbon concentrating mechanism, thereby reducing photorespiratory energy losses and allowing Rubisco to operate close to its maximum capacity. However, further enhancing C_4_ photosynthesis could have the potential for major agricultural impact. Maize, sorghum and sugarcane are among the most productive crops and all three belong to the NADP-ME biochemical subtype. Here, we used the closely related model NAPD-ME species *S.viridis* to study the effect of increased electron transport capacity on C_4_ photosynthesis. Since cyt*b*_6_*f* controls electron transport in C_3_ plants and its abundance is likely determined by the abundance of Rieske FeS subunit^6, 19^, we attempted to accelerate electron transport rate in *S.viridis* by overexpressing Rieske FeS. As shown by immunodetection of the thylakoid protein complexes isolated from mesophyll and bundle sheath cells, constitutive overexpression of Rieske FeS protein in *S.viridis* resulted in higher abundance of the complete cyt*b*_6_*f* complex in both cell types (Fig. 2).

### Cytochrome *b*_6_*f* complex regulates electron transport rate of C_4_ photosynthesis

The role of cyt*b*_6_*f* in the regulation of electron transport is defined by its position between the two photosystems. PSII activity can be controlled by cyt*b*_6_*f* levels either by limiting the rate of plastoquinol oxidation and hence oxidation of the primary electron acceptor Q_A_ or by down-regulation of photochemical efficiency via NPQ. Plants generated here with more cyt*b*_6_*f* had higher effective quantum yields of PSII at irradiances above 200 μmol m^−2^ s^−1^ which indicated that a higher portion of absorbed light reached PSII reaction centres (Fig. 4e). Lower NPQ measured in plants with higher cyt*b*_6_*f* abundance at those irradiances suggested that the increase in φPSII was attributed to the lower loss of energy via thermal dissipation (Fig. 5a). Both observations suggested that in FeS-OE plants Q_A_ was more oxidized and linear electron flow was less limited by the rate of plastoquinol oxidation. Since in NADP-ME monocots mesophyll cells contain a much higher portion of PSII than bundle sheath cells (95-98%^29^), leaf level changes in PSII light-use efficiency can be attributed predominantly to mesophyll cells suggesting a higher capacity for whole chain electron transport in mesophyll cells. Steady-state electron flow measurements in leaves under the growth irradiance also confirmed the increased capacity for linear electron flow in FeS-OE plants (Table 1).

PSI does not have an effective repair mechanism, such as the D1 protein turnover process of PSII^30^, and when the transfer of electrons to PSI exceeds the capacity of the stromal acceptors, PSI photoinhibition causes long-term inhibitory effects on electron transport and carbon fixation^31, 32^. Cyt*b*_6_*f* determines the amount of electrons reaching PSI from the intersystem chain, especially at low irradiance when cyclic electron flow (CEF) is mostly inactive^33^. In line with that, at irradiances below 340 μmol m^−2^ s^−1^ higher abundance of cyt*b*_6_*f* increased electron supply to the donor side of PSI (Fig. 5c) which resulted in the higher effective quantum yield of PSI (Fig. 5b).

Calculations of electron transport rates through PSI and PSII are complicated in C_4_ plants because of more complex distribution of light energy between two PSI and PSII in the two-cell system. However, we attribute more efficient use of absorbed light for photochemical reactions in both photosystems to a “release” of cyt*b*_6_*f* control in both mesophyll and bundle sheath cells. In C_3_ *A.thaliana* plants overexpressing Rieske FeS, increased φPSI and φPSII were also accompanied by increased abundances of all electron transport complexes^6^. In FeS-OE plants generated here, due to unaltered abundances of other photosynthetic complexes (Fig. 2, Fig. 7), higher φPSI and φPSII clearly demonstrate the effect of the cyt*b*_6_*f* control on whole leaf electron transport.

### Cyclic electron flow is regulated by Cytochrome *b*_6_*f*

In C_3_ plants, the rate of CEF around PSI increases at high light and generates a higher ΔpH which consequently decreases light harvesting efficiency via generation of a higher NPQ^34^. In line with CEF increasing electron supply to the donor side of PSI^35^, no difference in electron flux on the donor side of PSI was detected in plants with higher cyt*b*_6_*f* abundance at higher irradiance (Fig. 5c). Instead, a higher yield of PSI detected at higher irradiance corresponded to the increased availability of PSI electron acceptors (Fig. 5d). It has been suggested that heme *x* of cyt*b*_6_*f* exposed to stroma may be involved in the PGR5-mediated CEF route^36^ and cyt*b*_6_*f*, PSI and PGR5 can form a super-complex together with ferredoxin, ferredoxin:NADPH oxidoreductase and PROTON GRADIENT REGULATION5-like1^37^. If this is the case, higher abundance of cytb6f could directly increase the rate PGR5-mediated CEF and contribute to alleviating PSI acceptor side limitation. During the C_3_-C_4_ evolutionary transition increased PGR5 abundance appears to have been favoured in both cell types^38^ and therefore stimulatory effect of Rieske FeS overexpression on PGR5-mediated CEF could be valid for both mesophyll and bundle sheath cells.

C_4_ plants typically grow in high light environments which allow them to accommodate the higher ATP requirements of the pathway^39^. Higher *pmf* generated by the plants with increased cyt*b*_6_*f* abundance at high irradiance (Fig. 5e, Fig. 5g), conceivably, indicated a higher capacity for ATP production and could underpin higher assimilation rates (Fig. 4d). Since the effect of cyt*b*_6_*f* abundance on *pmf* was most prominent above 825 μmol m^−2^ s^−1^, this irradiance might be a prerequisite for a realisation of increased photosynthesis in C_4_ plants. Hence the plants with higher cyt*b*_6_*f* abundance grown here at the irradiance of 200 μmol m^−2^ s^−1^ did not have a growth advantage.

Since the electron transport chain in bundle sheath cells is specifically optimised for active CEF^27^, higher proton-motive force (*pmf*) and ΔpH detected in plants with higher cyt*b*_6_*f* abundance at high light (Fig. 5e, Fig. 5g) could be present in bundle sheath cells. The NDH-mediated route of CEF is considered to be prevalent in bundle sheath cells of NADP-ME plants^40^. Similar protein abundance of the NdhH subunit between FeS-OE and Null plants (Fig. 7) suggested that CEF activity in bundle sheath cells could be regulated also at the level of cyt*b*_6_*f* abundance. Lateral heterogeneity of the thylakoid membranes where PSII is localized to the stacked grana and PSI to the stromal lamellae^41^ provides the basis for functional specialization of cyt*b*_6_*f*. Since about 55% of cyt*b*_6_*f* is found in appressed grana and 45% are distributed over the stromal lamellae^42^, these two fractions of cyt*b*_6_*f* might be more specific for either linear or cyclic electron flow. In line with this we detected cyt*b*_6_*f*\*complex (Fig. 2) in bundle sheath cells of *S.viridis* that might represent a CEF-specific cyt*b*_6_*f* potentially forming a complex with its electron transport-partners.

### Effect of increased Cytochrome *b*_6_*f* levels on NPQ

Our results suggest that lower NPQ levels detected in plants with increased Cytochrome *b*_6_*f* levels underpinned more efficient use of light energy that could result in higher assimilation rates. Remarkably, higher abundance of Rieske FeS resulted in slower NPQ induction in both C_3_^6^ and C_4_ plants (Fig. 6) showing an apparent effect on pH-dependent qE component of NPQ. Moreover, here we demonstrate that FeS-OE plants generated the same or higher levels of *pmf* (and ΔpH) compared to Null plants despite the decreased NPQ (Fig. 5e).

Plants with higher cyt*b*_6_*f* levels retained linear relationships between *pmf* and NPQ, but showed an offset in the NPQ response (Fig. 8).

**Fig. 8.**
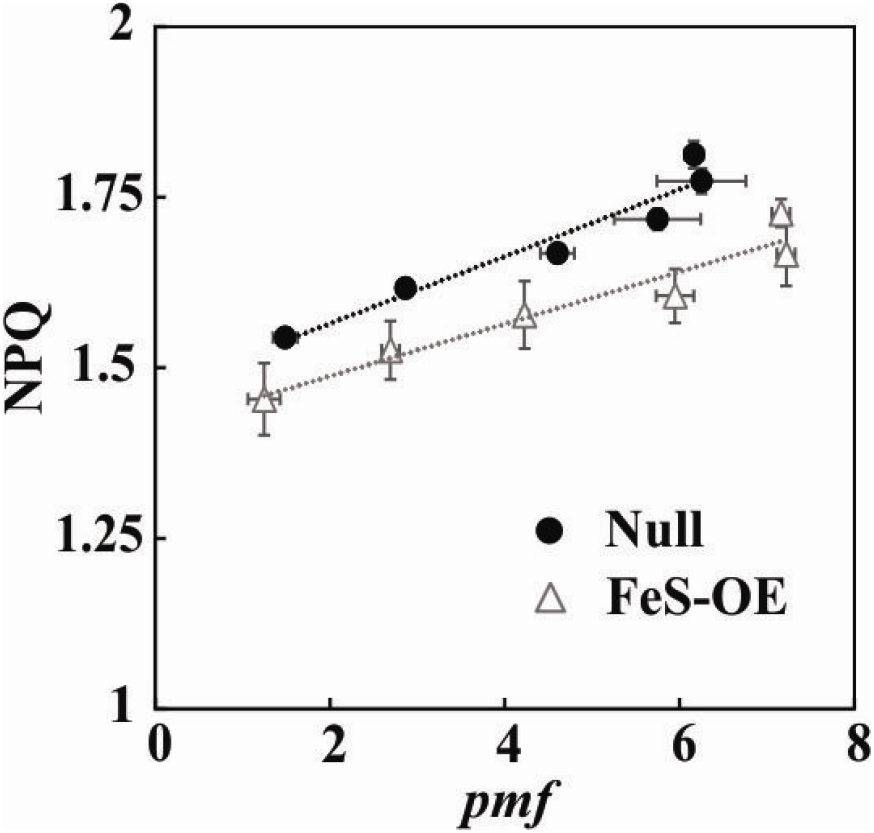
The relationship between light-induced proton-motive force (*pmf*) and nonphotochemical quenching (NPQ) in plants with Rieske FeS overexpression (FeS-OE) and Null plants. NPQ data from Fig. 5a were plotted against the steady-state *pmf* from Fig. 5e.

Although the regulation of ATP synthase is thought to play a key role in modulating *pmf*^43^ and NPQ^43^, the effect of Rieske FeS overexpression presented on Fig. 7 was not due to changes in proton conductivity (Fig. 5h). In addition, no decrease in violaxanthin content could be detected in *A.thaliana* plants overexpressing Rieske FeS^6^ and no difference in PsbS abundance was found between FeS-OE and Null plants in this work (Fig. 7). These observations suggest that there might be other mechanisms regulating NPQ in these plants. PGR5-mediated CEF might serve also for redox poising of the electron transport chain^44^ to prevent the reduction of Q_A_ and the closure of PSII centres. If higher cyt*b*_6_*f* abundance stimulated this pathway, it could result in a sustained change in the redox state of the Q_A_ pool (i.e. 10% more oxidised Q_A_) and hence a corresponding offset of NPQ response in FeS-OE plants (Fig. 8). While lower NPQ might be detrimental for C_3_ plants in high light conditions, due to higher availability of CO_2_ as terminal electron acceptor, C_4_ photosynthesis is more resistant to photo-damage by high irradiance^8^ and the “release” of cyt*b*_6_*f* control could have positive effects on photosynthesis without photoinhibitory effects on photosystems.

### C_4_ photosynthesis is co-limited by electron transport capacity

At low intercellular CO_2_ (C_i_), the assimilation rate is largely limited by the rate of primary CO_2_ fixation in mesophyll cells^45^, whilst at high C_i_, according to the C_4_ photosynthesis model^46, 47^, assimilation rate is co-limited by the amount and activity of Rubisco, the capacity for regeneration of either PEP or ribulose bisphosphate and the rate of whole leaf electron transport (the sum of electron transport rates of the two cell types). In line with the model, saturating rates of CO_2_ assimilation were about 10% higher in plants with about 10% higher Rieske FeS abundance (and hence with higher electron transport rate as discussed above). When measured in the steady-state, there was a strong positive correlation between Rieske FeS abundance and CO_2_ assimilation rate at ambient C_i_ in three independent transgenic lines (Fig. 3).

While very little work has been done on improving C_4_ photosynthesis by genetic modification, maize plants overexpressing both subunits of Rubisco together with the chaperonin RUBISCO ASSEMBLY FACTOR 1 demonstrated higher assimilation rates at high C_i_ and high light^48^. Importantly, since plants with higher cyt*b*_6_*f* abundance examined here did not have higher Rubisco content, both abundance of Rubisco and cyt*b*_6_*f* are likely co-limiting factors of C_4_ photosynthesis under high light and non-limiting CO_2_, in accordance with the C_4_ model predictions.

## Conclusions

We provide evidence that cyt*b*_6_*f* exerts a high level of control over the rate of electron transport in both mesophyll and bundle sheath cells in the C_4_ model plant *S.viridis*. We have demonstrated that increasing cyt*b*_6_*f* content can help improve light conversion efficiency in both photosystems and support generation of higher proton-motive force. Our results support the idea that C_4_ photosynthesis at ambient and high CO_2_ and high irradiance is co-limited by the electron transport rate. Engineering C_4_ crops with higher cyt*b*_6_*f* abundance may result in higher CO_2_ assimilation rates and higher yield.

## Materials and methods

### Construct generation

Two constructs, 230 and 231, were created using Golden Gate MoClo Plant Parts Kit^49^ (Fig. S1). The Golden Gate cloning system allows assembly of up to seven independent transcription units (modules 1-7) in a plant binary vector pAGM4723. In both constructs the first expression module was occupied by the hygromycin phosphotransferase gene (*hpt*) driven by the rice actin promoter (pOsAct). The second module in both constructs was taken by the coding sequence of PetC gene encoding for the Rieske FeS subunit of the cytochrome *b*6*f* complex from *Brachypodium distachyon* domesticated for the Golden Gate (Fig. S1) and driven by the maize ubiquitin promoter (p*Zm*Ubi). Construct 231 contained a third expression module with the coding sequence of the cyan fluorescent protein mTurquoise driven by the 2×35S promoter. The bacterial terminator tNos was used in all transcription modules. Both constructs were verified by sequencing and transformed into *Agrobacterium tumefaciens* strain AGL1 to proceed with stable plant transformation.

### Generation and selection of transgenic plants

Stable agrobacterium-mediated transformation of *S.viridis* (accession A10.1) tissue culture was performed as described in Osborn et al.^50^. Hygromycin-resistant plants were transferred to soil and grown in control environmental chambers with 2% CO_2_, the irradiance of 200 μmol m^−2^ s^−1^, 16 h photoperiod, 28 °C day, 20 °C night and 60% humidity. T_0_ plants were analysed for the *hpt* gene copy number by digital droplet PCR (ddPCR, iDNA genetics, UK) as described in Osborn et al.^50^. The plants that went through the transformation process and tested negative for the *hpt* gene were used as control for T_0_ and T_1_ generations. Null segregates were used as control for T_2_ plants. T_1_ and T_2_ progenies were analysed with ddPCR to confirm the presence of insertions. MultispeQ v1.0 leaf photosynthesis V1.0 protocol was used for a fast screening of transgenic plants for altered photosynthetic properties^51^. MultispeQ data were analysed with the PhotosynQ web application (https://photosynq.org).

### Plant growth conditions

The T_1_ progenies of T_0_ plants were analysed to select the lines with Rieske FeS overexpression. T_1_ seeds were sterilised and germinated on medium containing 2.15 g L^−1^ Murashige and Skoog ^17^ salts, 10 mL L^−1^ 100x MS vitamins stock, 30 g L^−1^ sucrose, 7 g L^−1^ Phytoblend, 20 mg L^−1^ hygromycin (pH 5.7). Seedlings that were able to develop secondary roots were transferred to 1 L pots with garden soil mix fertilised with 1 g L^−1^ Osmocote (Scotts, Bella Vista, Australia). Plants were grown in controlled environmental chambers under the irradiance of 200 μmol m^−2^ s^−1^, 16 h photoperiod, 28 °C day, 20 °C night, 60% humidity and ambient CO_2_ concentrations. Seeds of the T_1_ plants with confirmed Rieske FeS overexpression were collected to be analysed in the T_2_ generation. T_2_ seeds were incubated in 5% liquid smoke (Wrights, B&G foods, Parsippany, NJ) overnight and germinated in 1 L pots with garden soil mix layered on top with 2 cm seed raising mix (Debco, Tyabb, Australia) both containing 1 g L^−1^ Osmocote. Youngest fully expanded leaves of the 3-4 weeks plants were used for all analyses.

### Gas exchange measurements

Rates of CO_2_ assimilation were measured over a range of intercellular CO_2_ partial pressure and irradiances simultaneously with chlorophyll fluorescence using a portable gas-exchange system LI-6800 (LI-COR Biosciences, Lincoln, NE) and a Fluorometer head 6800-01A (LI-COR Biosciences). Leaves were first equilibrated at 400 ppm CO_2_ in the reference side, an irradiance of 1500 μmol m^−2^ s^−1^, leaf temperature 28 °C, 60% humidity and flow rate 300 μmol s^−1^. CO_2_ response curves were conducted under constant irradiance of 1500 μmol photons m^−2^ s^−1^ by imposing a stepwise increase of CO_2_ concentrations from 0 to 1600 ppm at 3 min intervals. Light response curves were measured at constant CO_2_ partial pressure of 400 ppm in the reference cell under a stepwise increase of irradiance from 0 to 3000 μmol m^−2^ s^−1^ at 2 min intervals. Red-blue actinic light (90%/10%) was used in all measurements.

### Protein isolation and Western blotting

To isolate proteins from leaves, leaf discs of 0.71 cm^2^ were collected and frozen immediately in liquid N_2_. One disc was ground in ice-cold glass homogeniser in 0.5 mL of protein extraction buffer: 100 mM trisaminomethane-HCl, pH 7.8, supplemented with 25 mM NaCl, 20 mM ethylenediaminetetraacetic acid, 2% sodium dodecyl sulfate (w/v), 10 mM dithiothreitol and 2% (v/v) protease inhibitor cocktail (Sigma, St Louis, MO). Protein extracts were incubated at 65 °C for 10 min and then centrifuged at 13000 g for 1 min at 4 °C to obtain clear supernatant. Protein extracts were supplemented with 4x XT Sample buffer (BioRad, Hercules, CA), loaded on leaf area basis and separated by polyacrylamide gel electrophoresis (Nu-PAGE 4-12% Bis-Tris gel, Invitrogen, Life Technologies Corporation, Carlsbad, CA). Proteins were transferred to a nitrocellulose membrane and probed with antibodies against various photosynthetic proteins purchased from Agrisera (Vännäs, Sweden). Quantification of immunoblots was performed with Image Lab software (Biorad, Hercules, CA).

### Thylakoid isolation and Blue Native gel electrophoresis

These procedures were done in dim light and at 4 °C (or on ice) to reduce light-induced damage of isolated thylakoid complexes. Ten youngest fully expanded leaves were collected from one plant and midribs were removed. Leaves were cut into 3 mm pieces and ground in 100 mL of ice-cold grinding buffer (50 mM Hepes-NaOH, pH 7.5, 330 mM sorbitol, 5 mM MgCl_2_) in Omni Mixer (Thermo Fisher Scientific, Tewksbury, MA) at the intensity #10 for 2 s. This fast treatment was used to break only mesophyll cells that don’t have suberised cell walls. The homogenate was passed through the 80-μm nylon filter and the filtrate containing mesophyll suspension was collected. All tissues collected on the filter were again homogenised in 100 mL of grinding buffer in Omni Mixer during three 10-s cycles at the intensity #7. The homogenate was passed through a tea strainer and then bundle sheath strands from the filtrate were collected on the 80-μm nylon filter. Bundle sheath strands were further ground in 10 mL of grinding buffer in ice-cold glass homogeniser.

Mesophyll and bundle sheath suspensions were filtered through a layer of Miracloth (Merck Millipore, Burlingtone, MA) and centrifuged at 6000 rpm, 4 °C for 5 min. Pellets were first resuspended in ice-cold shock buffer (50 mM Hepes-NaOH, pH 7.5, 5 mM MgCl_2_) and centrifuged again. Second time pellets were resuspended in ice-cold storage buffer (50 mM Hepes-NaOH, pH7.5, 100 mM sorbitol, 10 mM MgCl_2_) and centrifuged again. Finally pellets were resuspended in an equal aliquot of the storage buffer, snap-frozen in liquid N_2_ and stored at −80 °C.

For Blue Native gel electrophoresis - aliquots of thylakoid samples containing 10 μg chlorophyll (a+b) were taken for solubilisation. Aliquots were centrifuged at 6000 rpm, 4 °C for 5 min and then thylakoids were resuspended in ice-cold sample buffer (25 mM BisTris-HCl, pH 7.0, 20% (w/v) glycerol, 2.5% (v/v) protease inhibitor cocktail) to obtain chlorophyll (a+b) concentration of 1 μg μL^−1^. An equal volume of 2% (w/v) of n-dodecyl β-D-maltoside in sample buffer was added to thylakoids and the mixture was incubated for 5 min in darkness at 4 °C. Traces of unsolubilised material were removed by centrifugation at 14000 g at 4 °C for 30 min. Supernatants were supplemented with 10% (v/v) of Serve Blue G buffer (100 mM BisTris-HCl, pH 7.0, 0.5 M e-amino-n-caproic acid, 30% (w/v) sucrose, 50 mg mL^−1^ Serva Blue G) and applied to the precast native 3-12% Bis-Tris polyacrylamide gel (Invitrogen, Life Technologies Corporation, Carlsbad, CA). Blue Native gel electrophoresis was run according to Rantala et al.^52^. Ready gel was scanned, then incubated for 30 min in transfer buffer (25 mM trisaminomethane, 25 mM glycine, 20% methanol, 0.1% sodium dodecyl sulphate) and blotted to a nitrocellulose membrane. Western blotting was then performed as usually.

### Chlorophyll and Rubisco assays

For chlorophyll analysis frozen leaf discs were ground using the Qiagen TissueLyser II (Qiagen, Venlo, The Netherlands) and total chlorophyll was extracted in 80% acetone buffered with 25 mM HEPES-KOH. Chlorophyll *a* and *b* contents were measured at 750 nm, 663.3 nm and 646.6 nm, and calculated according to Porra et al.^53^. Amount of Rubisco active sites were estimated by [^14^C] carboxyarabinitol bisphosphate binding assay as described in Ruuska et al.^22^.

### Electron transport and electrochromic shift measurements

Effective quantum yield of PSII (φPSII)^54^ was probed simultaneously with the gas-exchange measurements under red-blue actinic light (90%/10%) using multiphase saturating flash of 8000 μmol m^−2^ s^−1^. Non-photochemical quenching (NPQ) as well as effective yields of photochemical and non-photochemical reactions in PSI were measured with Dual-PAM-100 (Heinz Walz, Effeltrich, Germany) under red actinic light with 300-ms saturating pulses of 10000 μmol m^−2^ s^−1^. The effective quantum yield of PSI (φPSI), the non-photochemical yield of PSI caused by donor side limitation (φND) and the non-photochemical yield of PSI caused by acceptor side limitation of PS I (φNA) were calculated as described earlier^55^. To monitor light-induced response of photosynthetic parameters leaves were dark-adapted for 30 min to record F0 and FM, the minimal and maximal levels of fluorescence in the dark, respectively. Afterwards saturating pulse was given after pre-illumination with far-red light to record PM, the maximal level of P700 oxidation, and P_0_, the minimal P700 signal, after the pulse. Next, photosynthetic parameters were monitored by the saturating pulse application every 45 s, first for 8 min under actinic light of 220 μmol m^−2^ s^−1^ and afterwards for 5 min in darkness to record NPQ relaxation. After that photosynthetic parameters were assessed in the same leaf over a range of irradiances from 0 to 1287 μmol m^−2^ s^−1^ at 1 min intervals.

The electrochromic shift signal was monitored as the absorbance change at 515 nm using a Dual-PAM-100 equipped with a P515/535 emitter-detector module (Heinz Walz, Effeltrich, Germany). Leaves were first dark adapted for 10 min and the absorbance change induced by a single turnover flash (ECS_ST_) was measured. Dark-interval relaxation of electrochromic shift signal was recorded during 3 min in darkness after 3-min intervals of illumination with actinic light at a stepwise increasing irradiance from 0 to 1287 μmol m^−2^ s^−1^. Total proton-motive force (*pmf*) was estimated from the amplitude of the rapid decay of the electrochromic shift signal and slow relaxation of the signal showed the contribution of proton gradient (ΔpH) and the electrochemical gradient (ΔΨ) across the thylakoid membrane^56, 57^. *Pmf* levels and the magnitudes of ΔpH and ΔΨ were normalized against the ECSST. Proton conductivity (g_H_^+^) of the thylakoid membrane through the ATP synthase was calculated as an inverse of the time constant obtained by fitting the first-order electrochromic shift relaxation^58^.

### RNA isolation and quantitative real-time PCR (qPCR)

Frozen leaf discs (0.71 cm^2^) were ground using the Qiagen TissueLyser II and RNA was extracted using the RNeasy Plant Mini Kit (Qiagen, Venlo, The Netherlands). DNA from the samples was removed using the Ambion TURBO DNA free kit (Thermo Fisher Scientific, Tewksbury, MA), and RNA quantity and quality were determined using a NanoDrop (Thermo Fisher Scientific, Tewksbury, MA). 1 μg of RNA was reverse transcribed into cDNA using SuperScript™ III Reverse Transcriptase (Thermo Fisher Scientific, Tewksbury, MA). qPCR and melt curve analysis were performed on a Viia7 Real-time PCR system (Thermo Fisher Scientific, Tewksbury, MA) using the Power SYBR green PCR Master Mix (Thermo Fisher Scientific, Tewksbury, MA) according to the manufacturer’s instructions. Primer pairs to distinguish between the PetC gene transcript from *S.viridis* and *B.distachyon* were designed using Primer3 in Geneious R9.1.1 (https://www.geneious.com): GCTGGGCAACGACATCAAG and CAAAGGAACTTGTTCTCGGC for SvPetC, GGCTCCGGGAGCAACAC and CAAAGGAACTTGTTCTCGGC for *Bd*PetC. Relative fold change was calculated by the ΔΔCt method, using the geometric mean of the Ct values for three reference genes described in Osborn et al.^50^.

### Statistical analysis

One-way ANOVA was performed for electron transport parameters measured with MultispeQ using the PhotosynQ web application (https://photosynq.org). For other measurements the relationship between mean values for transgenic lines and Nulls were tested using the Student’s t-test. Significant differences (*P*<0.05) are marked with asterisks.

## Acknowledgements

We thank Soumi Bala for the help with [^14^C] carboxyarabinitol bisphosphate binding assays and Tegan Norley for the help with electron transport measurements. This research was supported by the Australian Research Council Centre of Excellence for Translational Photosynthesis (CE140100015).

## Author contributions

S.vC. and R.F. conceived the project, S.vC., R.F., C.R. and M.E. planned experiments, P.L.- C. assembled constructs, M.E. performed experiments, M.E. analysed data and wrote the manuscript with contribution of all authors.

## Competing interests

The authors declare no competing interests.

## Supplementary materials

**Fig. S1.**
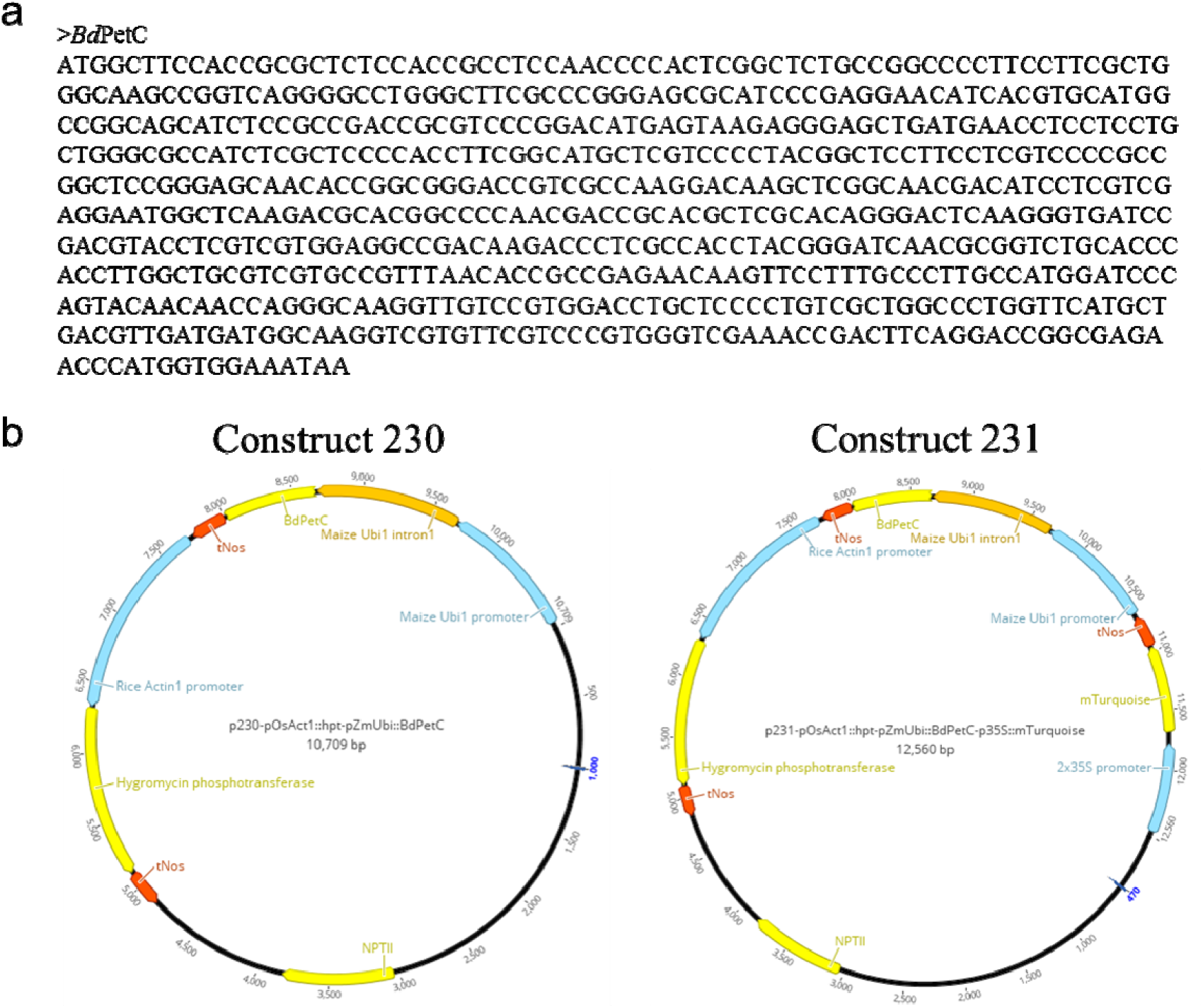
Constructs used for Rieske FeS overexpression in *S.viridis* **a.** Coding sequence of the PetC gene from *B.distachyon* (*Bd*PetC) domesticated for the Golden gate cloning system. **b.** Constructs for Rieske FeS overexpression containing the hygromycin phosphotransferase gene driven by the rice actin promoter and *Bd*PetC gene driven by the maize ubiquitin promoter. Construct 231 contained an additional expression module for expression of mTurquoise fluorescent protein under control of 2×35S promoter. The bacterial terminator tNos was used in all transcription modules.

**Fig. S2.**
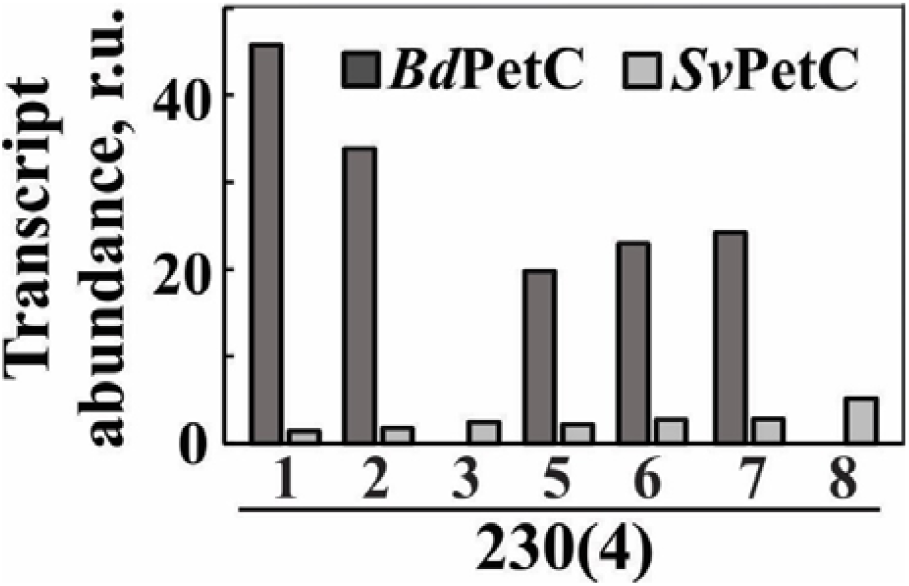
Transcript abundance of the PetC gene from *B.distachyon* (*Bd*PetC) or from *S.viridis* (*Sv*PetC) analysed in T_1_ plants of the line 230(4) relative to the expression level of the reference genes for Ubiquitin, Elongation factor 1a and Beta tubulin

**Fig. S3.**
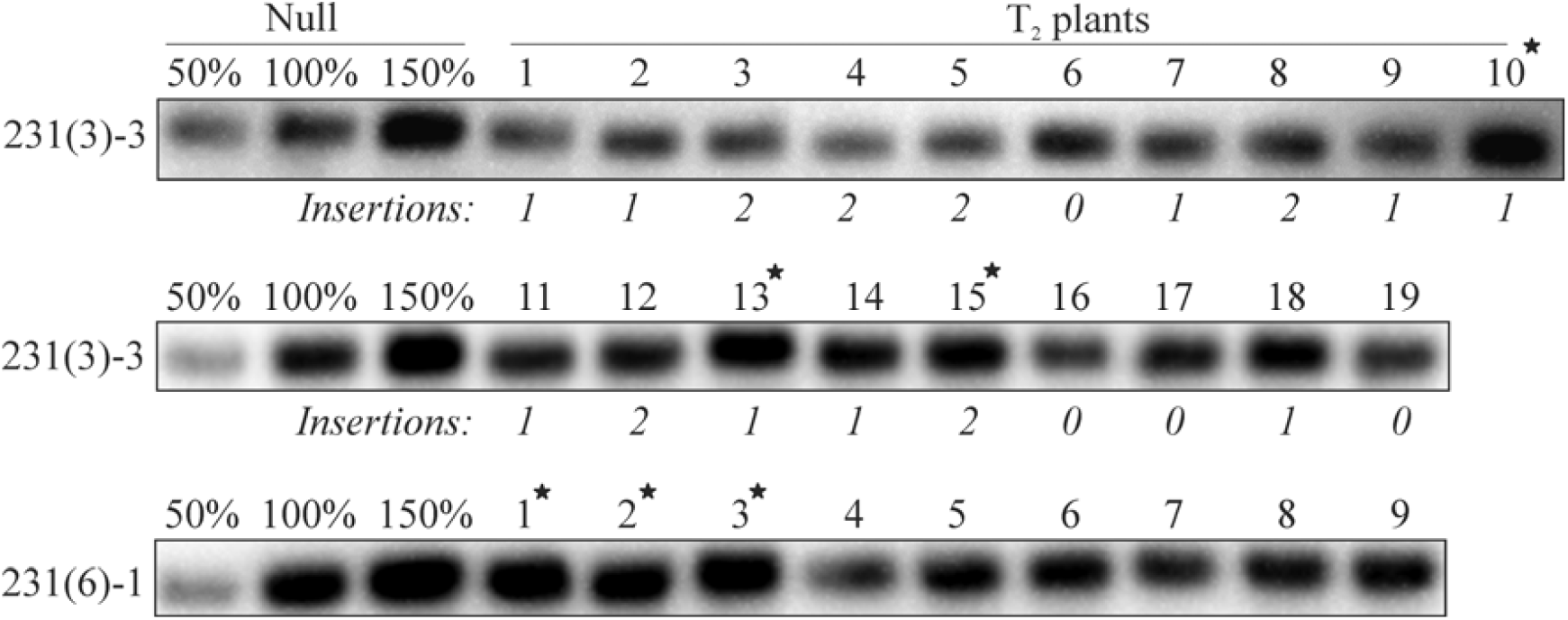
Immunodetection of Rieske FeS protein in T_2_ plants of lines 231(3)-3 and 231(6)-1 on leaf area basis. Insertion numbers indicate copy numbers of the hygromycin phosphotransferase gene; all T_2_ plants of line 231(6)-1 had 4 insertions. Asterisks indicate plants used for gas-exchange analysis on Fig. 3.

**Fig. S4.**
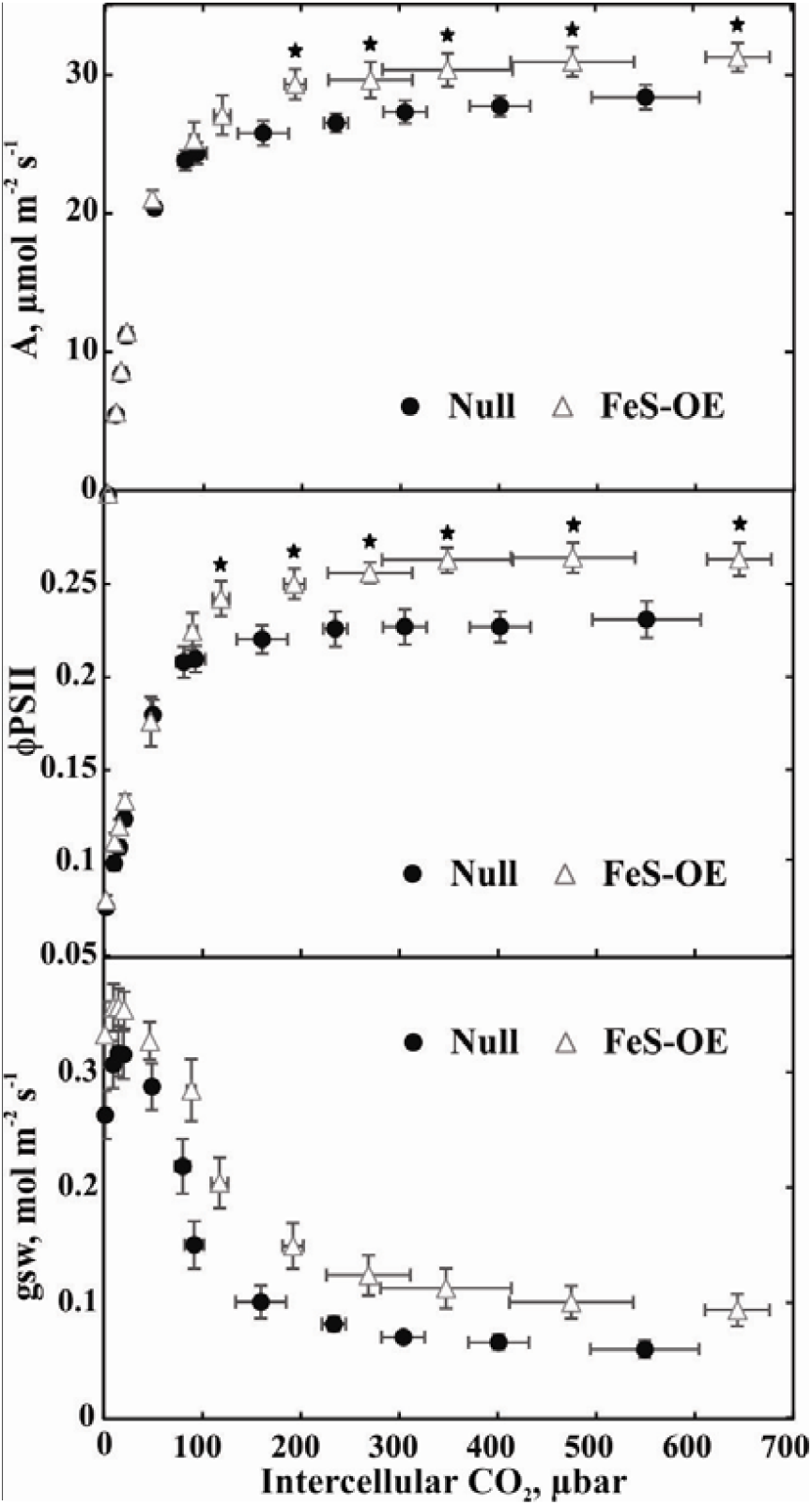
CO_2_ response of CO_2_ assimilation rate (A), quantum yield of Photosystem II (φPSII) and stomatal conductance (gsw) in T_2_ plants of line 231(3)-3 with Rieske FeS overexpression (FeS-OE) and Null plants measured at 1500 μmol m^−2^ s^−1^. Mean ± SE, n=3. Asterisks indicate statistically significant differences between transgenic and Null plants (*P*<0.05).

